# Draft Genome Assembly and Annotation of Red Raspberry *Rubus Idaeus*

**DOI:** 10.1101/546135

**Authors:** Haley Wight, Junhui Zhou, Muzi Li, Sridhar Hannenhalli, Stephen M. Mount, Zhongchi Liu

## Abstract

The red raspberry, *Rubus idaeus*, is widely distributed in all temperate regions of Europe, Asia, and North America and is a major commercial fruit valued for its taste, high antioxidant and vitamin content. However, *Rubus* breeding is a long and slow process hampered by limited genomic and molecular resources. Genomic resources such as a complete genome sequencing and transcriptome will be of exceptional value to improve research and breeding of this high value crop. Using a hybrid sequence assembly approach including data from both long and short sequence reads, we present the first assembly of the *Rubus idaeus* genome (Joan J. variety). The *de novo* assembled genome consists of 2,145 scaffolds with a genome completeness of 95.3% and an N50 score of 638 KB. Leveraging a linkage map, we anchored 80.1% of the genome onto seven chromosomes. Using over 1 billion paired-end RNAseq reads, we annotated 35,566 protein coding genes with a transcriptome completeness score of 97.2%. The *Rubus idaeus* genome provides an important new resource for researchers and breeders.

## Introduction

The red raspberry, *Rubus idaeus*, is widely distributed in all temperate regions of Europe, Asia, and North America and has been used as food and medicine since 4^th^ century AD (Graham et al., 2004). Often dubbed “European red raspberry”, *Rubus idaeus* is a globally commercialized specialty fruit crop with a large number of commercial varieties, high price, and increasing consumer demands. Owing to its health promoting value, unique flavor, and attractive appearance, *Rubus idaeus* sales have recently climbed by 8.4% with world production over 795, 000 tons (Darnell et al., 2006)(Barney et al., 2007; Food and Agriculture Organization of the United Nations Statistics Division (FAOSTAT)). In addition to its economic and health-promoting value, the red raspberry plants possess interesting and sometimes unique biological characteristics such as cold hardiness, aggregate fruits, perennial roots and biennial canes, either summer-bearing or ever-bearing flowering/fruiting, and large numbers of hybrids and cultivars. However, red raspberry breeding and research has fallen behind relative to other special fruit crops due to poor seed germination, an absence of reference genome, and limited transcriptome data (Graham and Woodhead, 2009; Hyun et al., 2014). With the recent publication of a high quality black raspberry (*Rubus occidentalis)* genome (VanBuren et al., 2018), this red raspberry genome allows comparative genomics, genetic breeding, and gene identification of this globally commercialized berry.

*Rubus idaeus* is a member of the economically important *Rosaceae* family that also includes rose, peach, apple, cherry, pear, almond, strawberry, and blackberry. Up to now, the genomes of several *Rosaceae* family members have been sequenced, including *Rubus occidentalis* (black raspberry) (VanBuren et al., 2016, 2018)*, Malus* x *domestica* (apple) (Daccord et al., 2017; Velasco et al., 2010), *Prunus persica* (peach) (Ahmad et al., 2011; Verde et al., 2013), *Pyrus bretschneideri* (Chinese pear) and *Pyrus communi* (Chagne et al., 2014) (Chagné et al., 2014), *Fragaria vesca* (woodland strawberry) (Edger et al., 2018; Shulaev et al., 2011), *Potentilla micranthia* (mock strawberry) (Buti et al., 2018), and *Rosa chinensis* (Chinese rose) (Hibrand Saint-Oyant et al., 2018; Raymond et al., 2018). Due to the small genome size, wide variety of fruit types (pomes, drupes, achenes, hips, follicles and capsules), and plant growth habits (ranging from herbaceous to cane, bush and tree forms), *Rosaceous* genomes offer one of the best systems for the comparative studies in genome evolution and development (Xiang et al., 2017). The availability of whole-genome sequences of key diploid species such as *Rubus idaeus* in this family will be crucial to these efforts.

Here, we report a draft genome assembly of red raspberry, *Rubus idaeus* (Joan J. variety), using long reads of single-molecular real-time (SMRT) Pacific Biosciences sequencing as well as high coverage Illumina short reads. The resulting draft genome is 300 Mbp in size with a BUSCO-calculated genome completeness score of 95.3% and contains 2,145 scaffolds with a N50 of 638 Kb. Using RNA-seq data from dissected fruit tissues at two developmental stages, we annotated the genome yielding 35,566 protein coding genes with a BUSCO-calculated transcriptome completeness score of 97.2%. We anchored the genome to two previously published high density linkage maps of *Rubus idaeus* (Ward et al., 2013), facilitating future marker development, breeding, and identification of genes controlling useful trait characteristics. Future comparative analysis, bolstered by this reference sequence, will enable the study of the complex evolution of morphological diversity in fleshy fruits of *Rosaceae*.

## Materials and methods

### Plant material and DNA sequencing

Joan J., a high-yielding, thornless, early primocane raspberry variety was chosen for genome sequencing. The Joan J. variety of *Rubus idaeus* was obtained from Appalachian Fruit Research Station of USDA ARS. Its genomic DNA was extracted from young leaves using the NucleoSpin® Plant II Midi kit (MACHEREY-NAGEL, Düren, Germany). DNA was sequenced at the Genomics Resource Center of the University of Maryland School of Medicine’s Institute of Genome Sciences. Specifically, a long read (5-20kb) PacBio genomic library was constructed using SMRTBell Template Prep Kit and sequenced on two SMRT cells on the PacBio Sequel System, generating 1,305,619 sequence reads with an average length of 9,879 bp (Supplementary Table 1). At the same time, a DNA-seq library was constructed using TruSeq DNA Library Pre Kits (Illumina) and then sequenced on Illumina HiSeq4000 platform in a single lane, yielding 249,081,860 reads of PE150 (Supplementary Table 1).

### Analysis of the Illumina DNA-Sequencing Data

PCR adapter sequences were removed using cutadapt (Martin, 2013). Jellyfish (Marçais and Kingsford, 2011) was then used to perform the k-mer distribution analysis with k=31 (Supplementary Figure 1).

### Genome Assembly

The genome assembly pipeline is shown in Supplementary Figure 2. A mixture of Illumina short reads and Pacbio long reads (Supplemental Table 1) were assembled into contigs using MaSuRCA; an assembler which combines the efficiency of de Bruijn graph and Overlap-Layout-Consensus approaches (Zimin et al., 2013). The specific settings used in the configuration file other than default were PE=pe 180 20, JF_SIZE = 200000000 and SOAP_ASSEMBLY=0. Subsequently, Redundans was used to remove heterozygous contigs using an all versus all BLAT approach (Pryszcz and Gabaldón, 2016). The Redundans pipeline also performed scaffolding using a mixture of Illumina short reads and Pacbio long reads (Pryszcz and Gabaldón, 2016). The genomes of *Potentilla micranthia* (Buti et al., 2018)*, Rubus occidentalis* (VanBuren et al., 2016), and *Fragaria vesca* (Edger et al., 2018) were leveraged to improve scaffolding using MeDuSa (Bosi et al., 2015). Scaffolds with less than 10X coverage were removed and scaffolds with more than 500 consecutive N’s were split. Bowtie2 version 2.3.0 (Langmead and Salzberg, 2012) was used to map the Illumina reads back onto the genome prior to Pilon with maximum fragment length to be 1000 and default settings otherwise. The mapping rate was 97.8% which further validates assembly quality. Pilon (Walker et al., 2014) was then used for one iteration to correct bases, fix misassembly and fill assembly gaps using the diploid parameter. Repeats were then softmasked by first creating a custom repeat library with RepeatModeler-1.0.11 (http://www.repeatmasker.org/RepeatModeler/) using the NCBI engine option and then using RepeatMasker (http://www.repeatmasker.org). Lastly, Haplomerger2 (Huang et al., 2017) split the resulting assembly into two sub-assemblies to further remove hetereozygosity.

### Sample collection and RNA-sequencing

Raspberry fruit from the Joan J. variety was dissected and separated into three tissues: ovary wall, seed (or ovule), and receptacle. The fruit was collected at two developmental stages, 0 and 12 DPA. Three biological replicates for above 6 tissues were obtained (Supplementary Table 2). Each tissue was homogenized in the presence of liquid nitrogen. Total RNA extraction was performed following a previously published protocol (Jones et al., 1997) with few modifications. The CTAB solution (3% CTAB, 100 mM Tris-HCl pH 8.0, 1.5 M NaCl, 20 mM EDTA, 5%PVP, and 1% β-mercaptoethanol made just before use) was added. 10 M Licl solution was mixed with total RNA for two days to precipitate RNA. The total RNA samples were eluted in DEPC-treated H_2_O and stored in −80°C.

Total RNA was shipped to the Weill Cornell’s Genomics and Epigenemics Core Facility, where polyA was isolated and RNA-seq libraries made using Tru-Seq RNA Library Prep Kit. Subsequently, the RNA-seq libraries were sequenced on Illumina HiSeq4000, yielding a total of 1,057,377,357 reads; an average of 96.24% of these reads mapped to the genome (Supplementary Table 2).

### Genome annotation

Repeat Masker was used with a custom repeat library built with Repeat Modeler to soft-mask the genome, and then a combination of *ab initio* and alignment guided assembly was employed to annotate the soft-masked genome. The Illumina Reads of RNA-seq data described above were trimmed with Trimmomatic (Bolger et al., 2014). RNA-Seq reads were mapped onto the draft genome sequence using Bowtie2 (Langmead et al., 2009). The bam file obtained was used to generate the training set for the gene prediction of BRAKER1 pipeline (Hoff et al., 2016). Candidate transcripts containing no known protein domains by Interproscan5 (Jones et al., 2014) were removed from the final set (13.96% percent decrease).

Trinity was then used to assemble the transcriptome on both genome guided and *de novo* settings (Grabherr et al., 2011). Prior to trinity assembly, reads were normalized using the perl script provided by Trinity and aligned using Bowtie2 (Grabherr et al., 2011; Langmead et al., 2009). Trinity assemblies were amassed into a comprehensive transcriptome database using PASA (Haas et al., 2003). Lastly, cd-hit-v4.6.8 (Li and Godzik, 2006) was used to cluster transcriptome assemblies from the resulting PASA and BRAKER1 assemblies with over 95% identity into unigenes. Unigenes that did not map to the genome, had no RNA-seq evidence, and had no known protein domains or orthologues were removed.

Blast2GOPro version 5.1.1 was used to associate Gene Ontology (GO) terms to the resulting transcripts (Supplementary Data 1). Protein sequences were searched against the non-redundant (nr) database protein database from NCBI using BLASTP with an e-value cutoff of 1.0E-3 (Conesa et al., 2005). InterProScan was run using default databases in order to assign putative domains to each transcript.

### GO enrichment

GO enrichment tests were performed to understand potential function of *Rubus* specific genes. GO term enrichment p-values were calculated using the Fisher’s exact test in the TopGO R package (http://bioconductor.org/packages/release/bioc/html/topGO.html). P-values were then adjusted using R’s FDR method.

### Anchorage to linkage maps

BLAT was run with default settings to identify unique and complete matches to each marker (Kent, 2002). After preparing the input files from BLAT (Supplementary Data 2), pseudochromosomes were then constructed using ALLMAPS with default parameters (Tang et al., 2015). Each genetic map was given a weight of 1. Chimeric scaffolds were manually broken at positions with low coverage, correcting many misassemblies. The seven pseudochromosomes were then constructed by integrating 98% of the markers from the genetic map.

### Comparative genomics

Orthology was established using OrthoFinder-1.1.2 (Emms and Kelly, 2015) using default parameters to infer a rooted species tree and identify orthologous gene groups. Subsequent to the gene trees Orthofinder also produced the species tree. The resulting orthogroups and species tree were then visualized with UpSetR (Conway et al., 2017) and an adjacent phylogenetic tree visualized with iTOL (Letunic and Bork, 2016) (Figure 2A). A Circos plot (Krzywinski et al., 2009) was created by creating links between every gene pair determined to be orthologs (Figure 2B-D). Syntenic orthologues were established by using MCScanX (Wang et al., 2012) with settings-s 5. An all by all BLASTp (Boratyn et al., 2013) query with an e-value cutoff of 1e-10 was performed and used as a basis for MCScanX with default parameters to identify syntenic gene regions.

## Results and Discussion

### Genome assembly and annotation

*Rubus idaeus* is a diploid species (2n=2x=14) with an estimated genome size of 293 Mbp based on flow cytometry analysis (Graham and Woodhead, 2009). We first sequenced the *Rubus idaeus* genome using 120X Illumina coverage (Supplementary Table 1). The distribution of k-mers indicates that the *Rubus idaeus* genome is approximately 303 Mbp (Methods), and the bimodal distribution of 31-mers (Supplemental Figure 1) suggests significant polymorphism and heterozygosity in the genome.

To overcome the issue of heterozygosity for genome assembly, a hybrid genome assembly approach was used taking advantage of both the sequencing depth and accuracy offered by the Illumina platform (at 120X coverage) and the sequence length offered by the PacBio platform (at 26X overage) (Supplemental Table 1). The pipeline of the assembly is outlined in Supplemental Figure 2. We used Redundans (Pryszcz and Gabaldón, 2016) and Haplomerger2 (Huang et al., 2017) tools to correct for heterozygosity. A comparative genomic approach (Bosi et al., 2015; Pop et al., 2004) was used as part of the genome assembly. Specifically, the most recently assembled genomes of closely related species *Potentilla micranthia* (Buti et al., 2018)*, Rubus occidentalis* (VanBuren et al., 2016), and *Fragaria vesca* (Edger et al., 2018) were leveraged to improve scaffolding using MeDuSa (Bosi et al., 2015). The resulting *R. idaeus* genome assembly is 300 Mbp in size, containing 2,145 scaffolds with a N50 of 638 Kb (Table 1). To assess the completeness of the genome, BUSCO v.3.0.2 (Simão et al., 2015) was used to locate the presence or absence of the embryophyta_odb9 (plant) dataset. The BUSCO Completeness Score reached 95.3% (Table 1), which validates the good assembly quality.

**Table 1.** Statistics of genome and transcriptome assemblies. Single (S), Duplicated (D), Fragmented (F) and Missing (M) single-copy orthologs are reported alongside the BUSCO completeness score.

|  |  |
| --- | --- |
| Total length | 300,259,977 bp |
| Scaffold N50 | 638,152 bp |
| Contig N50 | 250,294 bp |
| Smallest Scaffold | 501 bp |
| Largest Scaffold | 4,458,320 bp |
| N's | 174,429 bp (.000582%) |
| Sequence GC's | 37.9% |
| % Repeats | 43.35% |
| Busco Completeness Score (Genome) | 95.3% (S:86.1%, D:9.2%), F:1.5%, M:3.2% |
| Number of Annotated Protein Coding Genes | 35,566 |
| Busco Completeness Score (Transcriptome) | 97.2% (S:92.9%,D:4.3%), F:1.1%, M:1.7% |

To annotate the *Rubus idaeus* genome, a transcriptome was generated from 1,057,377,357 Illumina RNA-seq reads pooled from 18 RNA-seq libraries derived from three different fruit tissues (ovary wall, ovule/seed, receptacle) at two developmental stages (0 and 12 Days Post-Anthesis or DPA) in three biological replicates (Supplemental Table 2). A combination of *ab initio* and alignment guided assembly was employed to annotate the genome (soft-masked for repeats). This resulted in 35,566 protein coding genes with a BUSCO-calculated transcriptome completeness score of 97.2% (Table 1). The high completeness score indicates that transcripts from almost all genes expressed in these tissues have been sequenced. Finally, Blast2GO was used to associate Gene Ontology (GO) terms to the annotated genes (Supplementary Data 1).

### Anchoring scaffolds to genetic maps

The scaffolds were anchored onto pseudochromosomes (Figure 1) taking advantage of two previous genetic linkage maps. They are respectively the ‘Heritage’ and ‘Tulameen’ variety-based linkage maps that collectively contained 4225 markers. As a result, the pseudochromosomes contain 80.1% of the assembly (ie. at 240 Mb). The average magnitude of the Pearson correlation coefficient between the physical and map locations is 0.92 showing a high consistency between the genome and previously published linkage maps (Figure 1; Supplementary Data 2).

**Figure 1.**
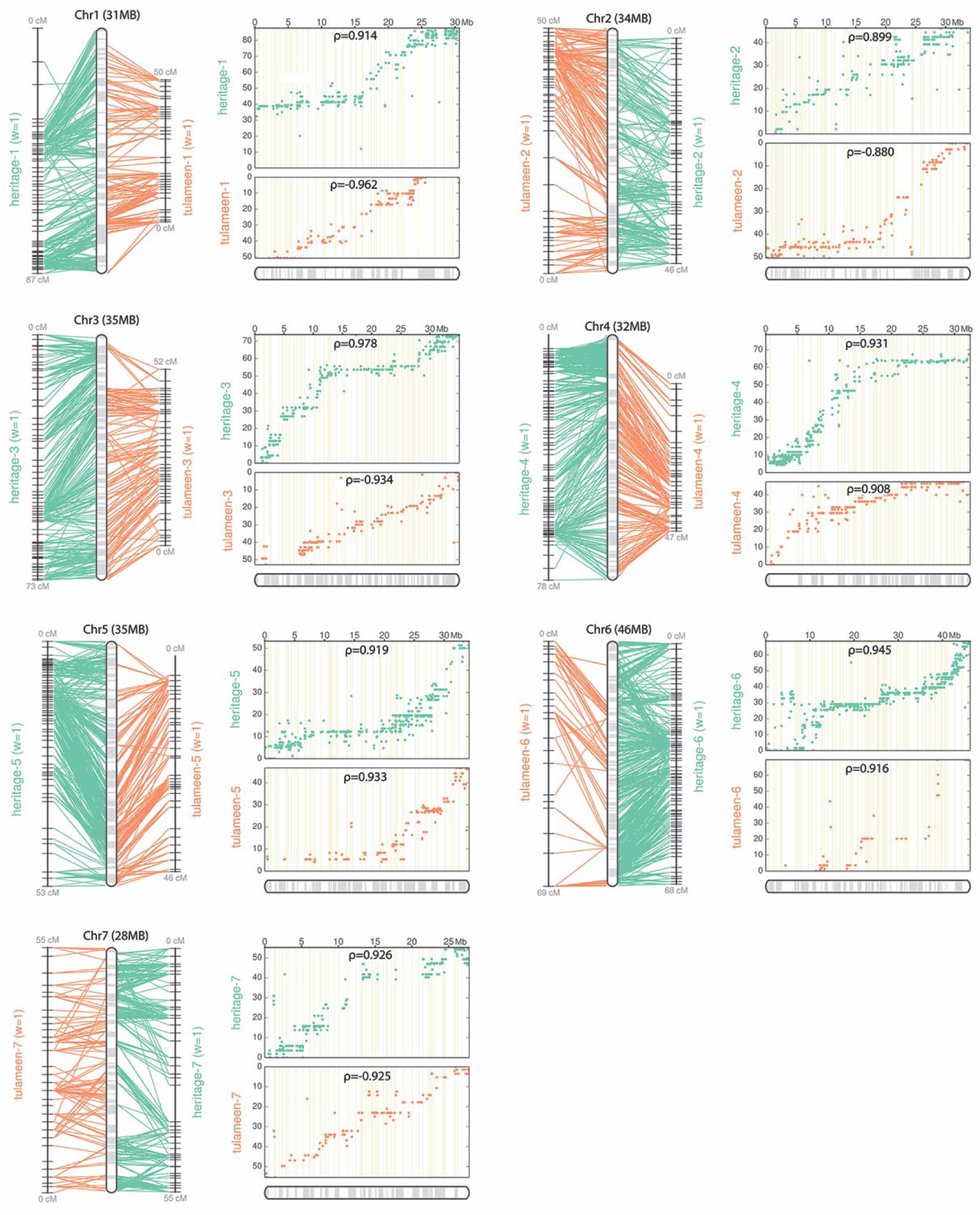
The correlation between physical map and the linkage maps of seven chromosomes. For each chromosome, the left figure illustrates the connections between physical positions on the assembled pseudomolecule and the two flanking linkage maps colored in orange and teal respectively. The orange coloring represents the tulmaneen linkage map whereas the teal represents the heritage linkage map (Ward et al., 2013). On the right is the scatter plot with dots representing the physical position on the chromosome (*x* axis) versus the map position (*y* axis). Rho (*ρ*) is the Pearson correlation coefficient (right panel). Each panel represents distinct chromosome.

### Comparative Genomics

Orthologous gene groups were established from 10 angiosperms using OrthoFinder-1.1.2 (Emms and Kelly, 2015); these include 9 members of the *Rosaceae* family (*Prunus persica*, *Pyrus communis*, *Malus* x *domestica*, *Rosa chinesis*, *Rosa multiflora*, *Rubus occidentalis*, *Rubus idaeus*, *Fragaria vesca*, *Potentilla micrantha*) and the model organism *Arabidopsis thaliana*, used here as an outlier species to root the tree. The resulting phylogenetic tree (Figure 2A) is consistent with previously published phylogenetic analyses of the *Rosaceae* family (Xiang et al., 2017). In total 25,193 orthogroups were established (Supplementary Data 3). As shown in Figure 2A, 10,205 orthogroups contained proteins from all 9 *Rosaceae* species as well as *Arabidopsis*. Interestingly, many specific orthogroups (1,878) are unique to *Malus* x *domestica* and *Pyrus communis*. Both species belong to the subfamily *Maleae*, which has undergone a whole genome duplication, at its origin (Daccord et al., 2017; Wu et al., 2013; Xiang et al., 2017). The large number of orthogroups shared between *Malus* x *domestica* and *Pyrus communis* suggests that substantial diversification occurred after whole genome duplication (WGD) within the *Maleae* subfamily, which may have contributed to the subfamily’s pome fruit type (Velasco et al., 2010; Xiang et al., 2017). Expectedly, all members of the *Rosaceae* family share many orthogroups (1,420) that are distinct from *Arabidopsis thaliana*. Members of the same genus also show a high number of common gene families. Specifically, there are 1,071 and 775 orthogroups limited to the *Rosa* and *Rubus* genera, respectively (Figure 2A, Supplementary Data 3). As *Rubus* is one of the largest and most morphologically diverse genus in the *Rosaceae* family (Alice and Campbell, 1999), we examined GO term enrichment among the 775 *Rubus*-specific orthogroups (Supplementary Data 4). Significantly enriched GO terms include chromatin assembly, RNA-splicing, and fungal-type cell wall organization, suggesting that *Rubus*-specific genes are involved in gene regulation and defense.

**Figure 2.**
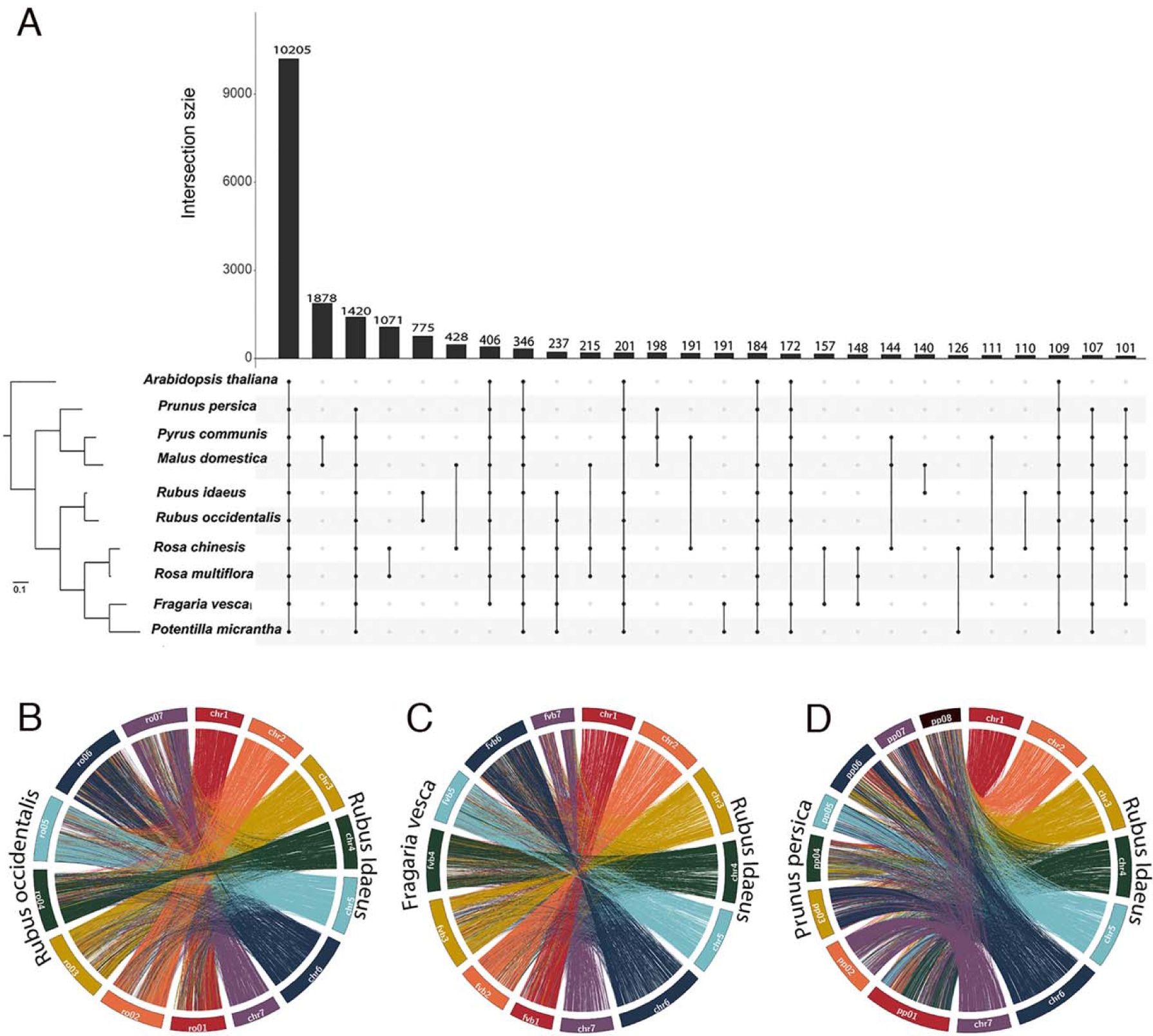
The distribution of shared gene families among nine *Rosaceae* species and *Arabidopsis thaliana*. (A) The left panel describes the phylogeny among the species. The branch length distances represent substitutions per site. The right panel is an UpSet plot (Conway et al., 2017): an alternative representation of a venn diagram with intersections (shared genes) greater than 100. The species described in each intersection is represented by the dotted lines, the size of the intersection is described by the bar chart above. (B) Circos plots (Krzywinski et al., 2009) displaying macrosynteny between the genomes of *Rubus idaeus* and *Rubus occidentalis*. (C) Macrosynteny between *Rubus idaeus* and *Fragaria vesca.* (D) Macrosynteny between *Rubus idaeus* and *Prunus persica*. For B to D, each connecting line represents an orthologous gene pair and the right half of each circle consists of the seven *Rubus ideaus* chromosomes colored by the spectral order in the rainbow.

Strawberry and raspberry share the same base chromosome number (n=7), with estimated divergence time of 75 million years (Xiang et al., 2017). *Rubus occidentalis* and *Rubus idaeus*, on the other hand, are closely related species. Syntenic blocks revealed a high collinearity between *Rubus idaeus* and *Rubus occidentalis* and between *Rubus idaeus* and *F. vesca* (Figure 2B and C). *R. occidentalis* had 25,289 gene pairs represented within 1,596 collinear blocks with *R. idaeus. F. vesca* and *R. idaeus* shared 17,769 syntenic gene pairs within 887 collinear blocks. This high degree of synteny helps validate the *Rubus idaeus* assembly. When compared with the more distant peach genome, *Prunus persica*, which has a different base chromosome number (n=8), collinearity decreases slightly: *P. persica* and *R. idaeus* share 17,064 gene pairs on 877 collinear regions. Although there is lower collinearity, there are strikingly large conserved syntenic blocks. For example, a large portion of *R. idaeus* chromosome 7 is syntenic to *P. persica* chromosome 2 while a smaller portion of *R. idaeus* chromosome 7 syntenic to *P. persica* 7 (Figure 2D).

To facilitate future functional studies of raspberry development, the *Rubus idaeus* genome assembly version 1 file, total transcript version 1 file, and annotation version 1 gff3 file are provided as Supplementary Data 5, 6, and 7 respectively. The Transcription Factors (TFs) and major hormonal pathway genes of *R. idaeus* are also identified and provided as Supplementary Data 8. Together with the GO assignment (Supplementary Data 1), linkage between physical and genetic markers (Supplementary Data 2), and ortholog assignment of nine *Rosaceae* species (Supplementary Data 3), these new genomic resources will assist raspberry research and breeding.

## Supporting information

terms to the annotated genes (Supplementary Data 1).

were established (Supplementary Data 3).

specific orthogroups (Supplementary Data 4).

Supplementary Data 5, 6, and 7 respectively

Supplementary Data 5, 6, and 7 respectively

Supplementary Data 5, 6, and 7 respectively

and provided as Supplementary Data 8.

## Supplemental Information

Supplementary Table 1. Summary statistics of DNA sequence data for *Rubus idaeus* genome assembly

Supplementary Table 2. Summary statistics of RNA-seq data for *Rubus idaeus* fruit tissues.

Supplementary Figure 1. Bimodal K-mer distribution of *Rubus idaeus* (variety Joan J.) genome.

Supplemental Figure 2. Genome assembly pipeline.

Supplementary Data 1: GO annotations associated with *Rubus idaeus* transcripts

Supplementary Data 2: Correlation between scaffold positions and genetic markers

Supplementary Data 3: Orthology clustering of *Rosaceae* species and *Arabidopsis*

Supplementary Data 4: GO enrichment of *Rubus*-specific genes

Supplementary Data 5: *Rubus idaeus*_genome_v1.fa.gz

Supplementary Data 6: *Rubus idaeus*_transcript_v1.fa.gz

Supplementary Data 7: *Rubus idaeus*_annotation_v1.gff3

Supplementary Data 8: Orthologs of known *Arabidopsis* transcription factors and hormone related genes

## Availability of supporting data

The genomic DNA-sequencing and RNA-sequencing data supporting the results of this article are available at Sequence Read Archive of NCBI with accession numbers SRP4284044 and SRP153061 respectively.

## Acknowledgements

We would like to thank Drs. Ann Callahan and Chris Dardick at USDA ARS for the Joan J. *Rubus idaeus* plants and Miss Anuhyea Pulapaka for help with the genome assembly.

## Supplemental Tables

**Supplementary Table 1.** Summary statistics of DNA sequence data for *Rubus idaeus* genome assembly

|  | **Mean Read length** | **Read count** | **Total base, bp** |
| --- | --- | --- | --- |
| Illumina PE | 150 | 249,081,860 | 37,455,877,274 |
| PacBio | 9,879 | 1,305,619 | 8,007,129,543 |

**Supplementary Table 2.** Summary statistics of RNA-seq data for *Rubus idaeus* fruit tissues.

| Sample | Number of Reads | % of Uniquely Mapped Reads | % of reads mapped to multiple loci | Total % reads mapped |
| --- | --- | --- | --- | --- |
| Ovule-0-26 | 60144925 | 87.94% | 7.13% | 95.07% |
| Ovule-0-41 | 57979785 | 89.56% | 7.24% | 96.80% |
| Ovule-0-7 | 66004490 | 89.29% | 7.30% | 96.59% |
| Receptacle-0-17 | 64293938 | 89.79% | 7.16% | 96.95% |
| Receptacle-0-27 | 61401941 | 89.54% | 7.18% | 96.72% |
| Receptacle-0-41 | 68480278 | 88.20% | 7.18% | 95.38% |
| Receptacle-12-13 | 67769659 | 89.39% | 6.32% | 95.71% |
| Receptacle-12-1 | 54088666 | 91.41% | 6.22% | 97.63% |
| Receptacle-12-4 | 50260693 | 90.69% | 6.70% | 97.39% |
| Seed-12-13 | 53332584 | 84.84% | 8.07% | 92.91% |
| Seed-12-1 | 55781661 | 89.18% | 8.47% | 97.65% |
| Seed-12-7 | 62967294 | 89.73% | 7.99% | 97.72% |
| Wall-0-24 | 65304690 | 89.32% | 7.27% | 96.59% |
| Wall-0-7 | 59200984 | 89.64% | 7.51% | 97.15% |
| Wall-0-13 | 71863354 | 89.68% | 7.34% | 97.02% |
| Wall-12-13 | 68284797 | 83.95% | 8.43% | 92.38% |
| Wall-12-1 | 70217618 | 88.17% | 8.27% | 96.44% |
| Wall-12-4 | 61592260 | 89.07% | 9.06% | 98.13% |

\*Sample names are “Tissue-Stage-unique ID of the specific sample”. The two stages are 0 DPA and 12 DPA. The tissues are Ovule, Seed, Receptacle, and Wall (ovary wall).

## Supplemental Figures

**Supplementary Figure 1.**
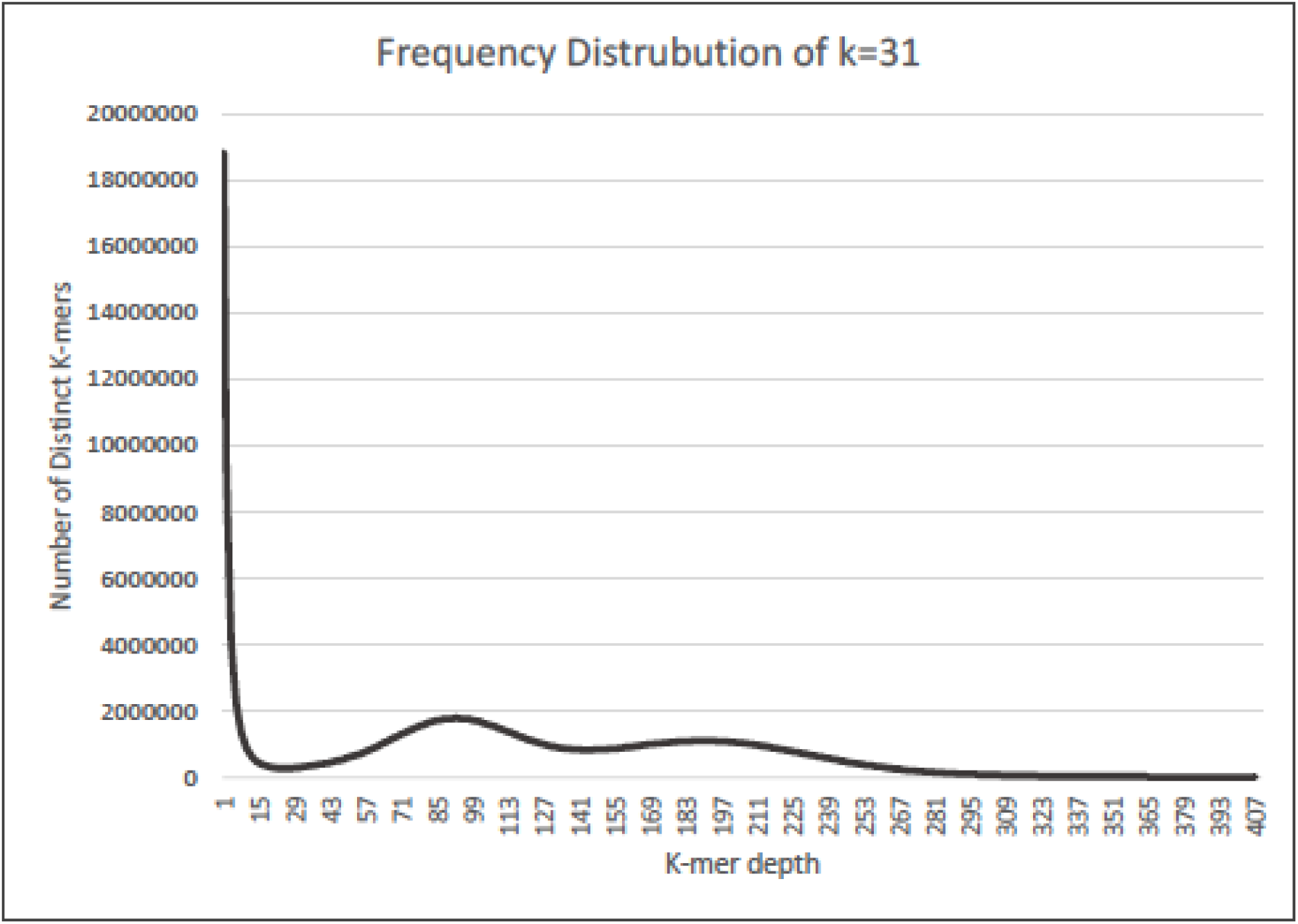
Bimodal K-mer distribution of *Rubus idaeus* (variety Joan J.) genome 31-mer distribution of *Rubus idaeus* genome obtained, using jellyfish, from 150-bp paired-end whole genome sequencing data.

**Supplemental Figure 2.**
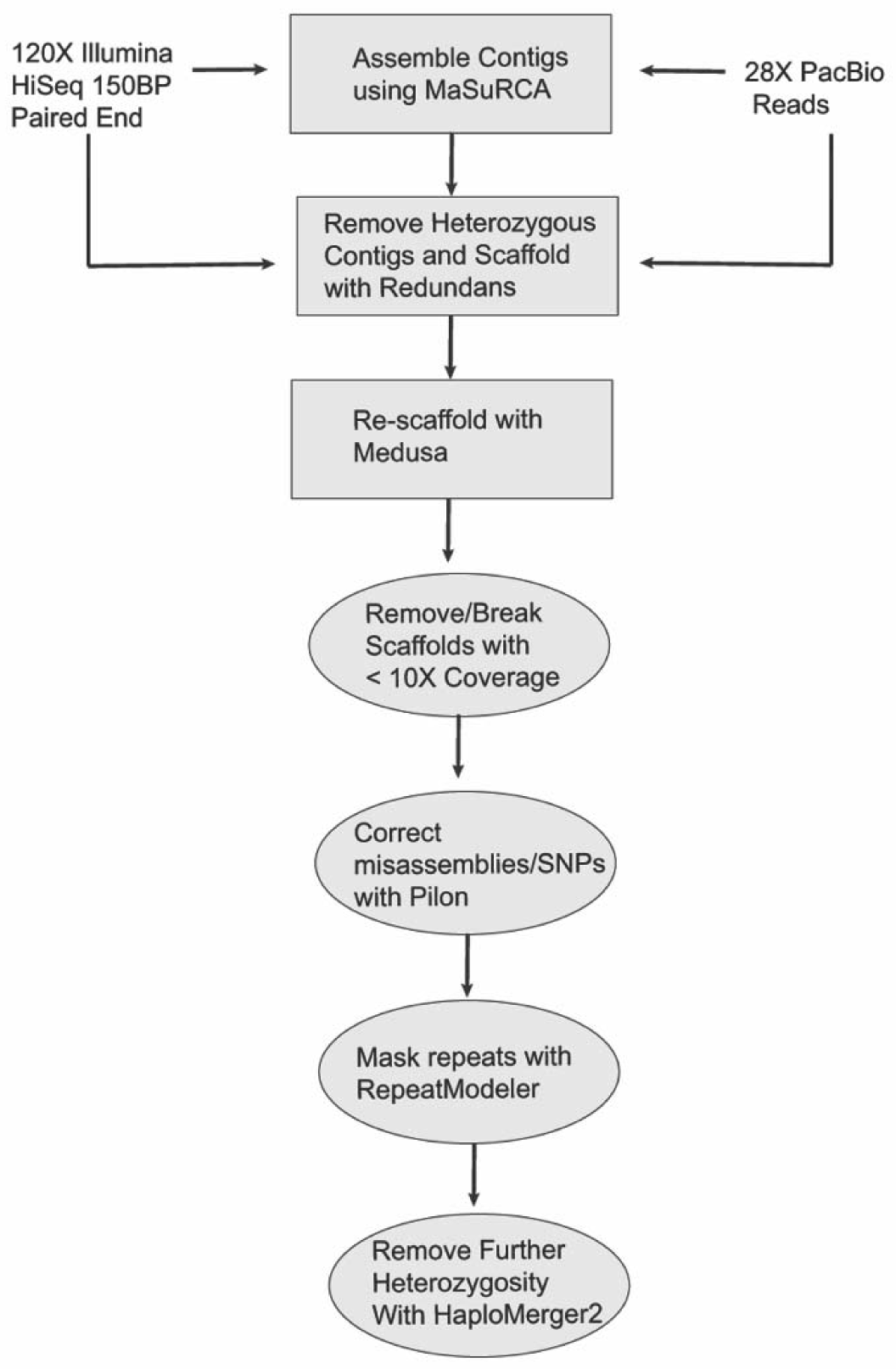
Genome assembly pipeline. Flowchart represents all steps of the genome assembly process upstream of anchoring to the linkage map.

## References

Ahmad, R., Parfitt, D.E., Fass, J., Ogundiwin, E., Dhingra, A., Gradziel, T.M., Lin, D., Joshi, N.A., Martinez-Garcia, P.J., and Crisosto, C.H. (2011). Whole genome sequencing of peach (Prunus persica L.) for SNP identification and selection. BMC Genomics 12, 569.

Alice, L.A., and Campbell, C.S. (1999). Phylogeny of Rubus (rosaceae) based on nuclear ribosomal DNA internal transcribed spacer region sequences. Am. J. Bot. 86, 81–97.

Barney, D.L., Bristow, P., Cogger, C., Fitzpatrick, S.M., Hart, J., Kaufman, D., Miles, C., Miller, T., Moore, P.P., Murray, T. and Rempel, H. (2007). Commercial red raspberry production in the Pacific Northwest. Pacific Northwest Ext. Publ. PNW, 598.

Bolger, A.M., Lohse, M., and Usadel, B. (2014). Trimmomatic: a flexible trimmer for Illumina sequence data. Bioinformatics (Oxford, England) 30, 2114–2120.

Boratyn, G.M., Camacho, C., Cooper, P.S., Coulouris, G., Fong, A., Ma, N., Madden, T.L., Matten, W.T., McGinnis, S.D., Merezhuk, Y., et al. (2013). BLAST: a more efficient report with usability improvements. Nucleic Acids Res 41, W29–W33.

Bosi, E., Donati, B., Galardini, M., Brunetti, S., Sagot, M.-F., Lió, P., Crescenzi, P., Fani, R., and Fondi, M. (2015). MeDuSa: a multi-draft based scaffolder. Bioinformatics 31, 2443–2451.

Buti, M., Moretto, M., Barghini, E., Mascagni, F., Natali, L., Brilli, M., Lomsadze, A., Sonego, P., Giongo, L., Alonge, M. and Velasco, R., (2018). The genome sequence and transcriptome of Potentilla micrantha and their comparison to Fragaria vesca (the woodland strawberry). GigaScience, 7(4), pp. 1–14.

Chagné, D., Crowhurst, R.N., Pindo, M., Thrimawithana, A., Deng, C., Ireland, H., Fiers, M., Dzierzon, H., Cestaro, A., Fontana, P., et al. (2014). The Draft Genome Sequence of European Pear (Pyrus communis L. “Bartlett”). PLoS ONE 9, e92644–e92644.

Conesa, A., Gotz, S., Garcia-Gomez, J.M., Terol, J., Talon, M., and Robles, M. (2005). Blast2GO: a universal tool for annotation, visualization and analysis in functional genomics research. Bioinformatics 21, 3674–3676.

Conway, J.R., Lex, A., and Gehlenborg, N. (2017). UpSetR: an R package for the visualization of intersecting sets and their properties. Bioinformatics 33, 2938–2940.

Daccord, N., Celton, J.-M., Linsmith, G., Becker, C., Choisne, N., Schijlen, E., van de Geest, H., Bianco, L., Micheletti, D., Velasco, R., et al. (2017). High-quality de novo assembly of the apple genome and methylome dynamics of early fruit development. Nature Genetics 49, 1099–1106.

Darnell, R.L., Alvarado, H.E., Williamson, J.G., Brunner, B., Plaza, M., and Negrón, E. (2006). Annual, Off-season Raspberry Production in Warm Season Climates. HortTechnology 16, 92–97.

Edger, P.P., VanBuren, R., Colle, M., Poorten, T.J., Wai, C.M., Niederhuth, C.E., Alger, E.I., Ou, S., Acharya, C.B., Wang, J., et al. (2018). Single-molecule sequencing and optical mapping yields an improved genome of woodland strawberry (Fragaria vesca) with chromosome-scale contiguity. GigaScience 7, 1–7.

Emms, D.M., and Kelly, S. (2015). OrthoFinder: solving fundamental biases in whole genome comparisons dramatically improves orthogroup inference accuracy. Genome Biology 16, 157.

Grabherr, M.G., Haas, B.J., Yassour, M., Levin, J.Z., Thompson, D.A., Amit, I., Adiconis, X., Fan, L., Raychowdhury, R., Zeng, Q., et al. (2011). Full-length transcriptome assembly from RNA-Seq data without a reference genome. Nature Biotechnology 29, 644–652.

Graham J., Woodhead M. (2009) Raspberries and Blackberries: The Genomics of Rubus. In: Folta K.M., Gardiner S.E. (eds) Genetics and Genomics of Rosaceae. Plant Genetics and Genomics: Crops and Models, vol 6. Springer, New York, NY

Graham, J., Smith, K., MacKenzie, K., Jorgenson, L., Hackett, C., and Powell, W. (2004). The construction of a genetic linkage map of red raspberry (Rubus idaeus subsp. idaeus) based on AFLPs, genomic-SSR and EST-SSR markers. Theor Appl Genet 109, 740–749.

Haas, B.J., Delcher, A.L., Mount, S.M., Wortman, J.R., Smith, R.K., Hannick, L.I., Maiti, R., Ronning, C.M., Rusch, D.B., Town, C.D., et al. (2003). Improving the Arabidopsis genome annotation using maximal transcript alignment assemblies. Nucleic Acids Research 31, 5654–5666.

Hibrand Saint-Oyant, L., Ruttink, T., Hamama, L., Kirov, I., Lakhwani, D., Zhou, N.N., Bourke, P.M., Daccord, N., Leus, L., Schulz, D., et al. (2018). A high-quality genome sequence of Rosa chinensis to elucidate ornamental traits. Nature Plants 4, 473–484.

Hoff, K.J., Lange, S., Lomsadze, A., Borodovsky, M., and Stanke, M. (2016). BRAKER1: Unsupervised RNA-Seq-Based Genome Annotation with GeneMark-ET and AUGUSTUS: Table 1. Bioinformatics 32, 767–769.

Huang, S., Kang, M., and Xu, A. (2017). HaploMerger2: rebuilding both haploid sub-assemblies from high-heterozygosity diploid genome assembly. Bioinformatics 33, 2577–2579.

Hyun, T.K., Lee, S., Rim, Y., Kumar, R., Han, X., Lee, S.Y., Lee, C.H., and Kim, J.-Y. (2014). De-novo RNA Sequencing and Metabolite Profiling to Identify Genes Involved in Anthocyanin Biosynthesis in Korean Black Raspberry (Rubus coreanus Miquel). PLoS One 9(2), e88292.

Jones, C.S., Iannetta, P.P.M., Woodhead, M., Davies, H.V., McNicol, R.J., and Taylor, M.A. (1997). The isolation of RNA from raspberry (Rubus idaeus) fruit. Molecular Biotechnology 8, 219–221.

Jones, P., Binns, D., Chang, H.-Y., Fraser, M., Li, W., McAnulla, C., McWilliam, H., Maslen, J., Mitchell, A., Nuka, G., et al. (2014). InterProScan 5: genome-scale protein function classification. Bioinformatics (Oxford, England) 30, 1236–1240.

Kent, W.J. (2002). BLAT--the BLAST-like alignment tool. Genome Research 12, 656–664.

Krzywinski, M., Schein, J., Birol, I., Connors, J., Gascoyne, R., Horsman, D., Jones, S.J., and Marra, M.A. (2009). Circos: an information aesthetic for comparative genomics. Genome Research 19, 1639–1645.

Langmead, B., and Salzberg, S.L. (2012). Fast gapped-read alignment with Bowtie 2. Nature Methods 9, 357–359.

Langmead, B., Trapnell, C., Pop, M., and Salzberg, S.L. (2009). Ultrafast and memory-efficient alignment of short DNA sequences to the human genome. Genome Biology 10, R25.

Letunic, I., and Bork, P. (2016). Interactive tree of life (iTOL) v3: an online tool for the display and annotation of phylogenetic and other trees. Nucleic Acids Res. 44, W242–245.

Li, W., and Godzik, A. (2006). Cd-hit: a fast program for clustering and comparing large sets of protein or nucleotide sequences. Bioinformatics 22, 1658–1659.

Marçais, G., and Kingsford, C. (2011). A fast, lock-free approach for efficient parallel counting of occurrences of k-mers. Bioinformatics (Oxford, England) 27, 764–770.

Martin, M. (2013). Cutadapt removes adapter sequences from high-throughput sequencing reads. EMBnet.journal 17, 10–12.

Pop, M., Kosack, D.S., and Salzberg, S.L. (2004). Hierarchical scaffolding with Bambus. Genome Research 14, 149–159.

Pryszcz, L.P., and Gabaldón, T. (2016). Redundans: an assembly pipeline for highly heterozygous genomes. Nucleic Acids Research 44, e113.

Raymond, O., Gouzy, J., Just, J., Badouin, H., Verdenaud, M., Lemainque, A., Vergne, P., Moja, S., Choisne, N., Pont, C., et al. (2018). The Rosa genome provides new insights into the domestication of modern roses. Nature Genetics 50, 772–777.

Shulaev, V., Sargent, D.J., Crowhurst, R.N., Mockler, T.C., Folkerts, O., Delcher, A.L., Jaiswal, P., Mockaitis, K., Liston, A., Mane, S.P., et al. (2011). The genome of woodland strawberry (Fragaria vesca). Nature Genetics 43, 109–116.

Simão, F.A., Waterhouse, R.M., Ioannidis, P., Kriventseva, E. V., and Zdobnov, E.M. (2015). BUSCO: assessing genome assembly and annotation completeness with single-copy orthologs. Bioinformatics 31, 3210–3212.

Tang, H., Zhang, X., Miao, C., Zhang, J., Ming, R., Schnable, J.C., Schnable, P.S., Lyons, E., and Lu, J. (2015). ALLMAPS: robust scaffold ordering based on multiple maps. Genome Biology 16, 3.

VanBuren, R., Bryant, D., Bushakra, J.M., Vining, K.J., Edger, P.P., Rowley, E.R., Priest, H.D., Michael, T.P., Lyons, E., Filichkin, S.A., et al. (2016). The genome of black raspberry (*Rubus occidentalis*). The Plant Journal 87, 535–547.

VanBuren, R., Wai, C.M., Colle, M., Wang, J., Sullivan, S., Bushakra, J.M., Liachko, I., Vining, K.J., Dossett, M., Finn, C.E., et al. (2018). A near complete, chromosome-scale assembly of the black raspberry (Rubus occidentalis) genome. Gigascience 7.

Velasco, R., Zharkikh, A., Affourtit, J., Dhingra, A., Cestaro, A., Kalyanaraman, A., Fontana, P., Bhatnagar, S.K., Troggio, M., Pruss, D., et al. (2010). The genome of the domesticated apple (Malus × domestica Borkh.). Nature Genetics 42, 833–839.

Verde, I., Abbott, A.G., Scalabrin, S., Jung, S., Shu, S., Marroni, F., Zhebentyayeva, T., Dettori, M.T., Grimwood, J., Cattonaro, F., et al. (2013). The high-quality draft genome of peach (Prunus persica) identifies unique patterns of genetic diversity, domestication and genome evolution. Nature Genetics 45, 487–494.

Walker, B.J., Abeel, T., Shea, T., Priest, M., Abouelliel, A., Sakthikumar, S., Cuomo, C.A., Zeng, Q., Wortman, J., Young, S.K., et al. (2014). Pilon: An Integrated Tool for Comprehensive Microbial Variant Detection and Genome Assembly Improvement. PLoS ONE 9, e112963.

Wang, Y., Tang, H., DeBarry, J.D., Tan, X., Li, J., Wang, X., Lee, T., Jin, H., Marler, B., Guo, H., et al. (2012). MCScanX: a toolkit for detection and evolutionary analysis of gene synteny and collinearity. Nucleic Acids Res 40, e49.

Ward, J.A., Bhangoo, J., Fernández-Fernández, F., Moore, P., Swanson, J., Viola, R., Velasco, R., Bassil, N., Weber, C.A., and Sargent, D.J. (2013). Saturated linkage map construction in Rubus idaeus using genotyping by sequencing and genome-independent imputation. BMC Genomics 14, 2.

Wu, J., Wang, Z., Shi, Z., Zhang, S., Ming, R., Zhu, S., Khan, M.A., Tao, S., Korban, S.S., Wang, H., et al. (2013). The genome of the pear (Pyrus bretschneideri Rehd.). Genome Research 23, 396–408.

Xiang, Y., Huang, C.-H., Hu, Y., Wen, J., Li, S., Yi, T., Chen, H., Xiang, J., and Ma, H. (2017). Evolution of Rosaceae Fruit Types Based on Nuclear Phylogeny in the Context of Geological Times and Genome Duplication. Molecular Biology and Evolution 34, 262–281.

Zimin, A. V., Marçais, G., Puiu, D., Roberts, M., Salzberg, S.L., and Yorke, J.A. (2013). The MaSuRCA genome assembler. Bioinformatics 29, 2669–2677.

